# Alpha-Beta oscillations implement inhibition of the ventral attention network during an attention task

**DOI:** 10.1101/2025.08.03.668152

**Authors:** M. Ferez, L. Luther, M. Bonnefond

## Abstract

Selective attention is a crucial mechanism that allows us to focus on relevant information while ignoring irrelevant information. Although extensive literature has proposed that alpha oscillations (7-14 Hz) suppress distractor processing, recent studies have questioned this role. In this magnetoencephalography (MEG) study, we used a modified Stroop task to investigate whether (1) alpha oscillations are associated with functional inhibition in higher-order visual regions and the ventral attention network (VAN), and (2) alpha phase adjusts in anticipation of relevant and irrelevant stimuli. We found no significant increase in pre-stimulus alpha power or phase adjustment over higher-order visual regions in the attend-color condition, consistent with the absence of a behavioral Stroop effect. However, we observed elevated alpha-beta power (10-20 Hz) in the VAN during both attend-color and attend-word conditions compared to control conditions, occurring before stimulus onset. Higher alpha-beta power in the right temporo-parietal junction correlated with faster reaction times, suggesting that inhibition of this network region facilitates task performance. This alpha-beta modulation in the VAN may represent a general mechanism for resisting distraction, preventing attentional capture by irrelevant information regardless of task condition. Additionally, we found enhanced theta power (4 Hz) over the VAN and left medial frontal gyrus (part of the cognitive control network) in both experimental conditions. Theta power correlated with improved reaction times across these regions. Furthermore, theta activity in the left medial frontal gyrus synchronized with VAN nodes, potentially indicating cross-network interaction between cognitive control and attention systems.

## Introduction

Selective attention is a crucial mechanism that enables us to focus on relevant information while ignoring irrelevant information. For example, the cocktail party effect shows that it is possible to listen to one person talking to you while ignoring other input streams of speech (Cherry, 1953), and ignoring elements of a stimulus in a working memory task has been shown to improve memory efficiency (Zanto & Gazzaley, 2009). Understanding this mechanism therefore seems essential.

Over the past few years, many research groups have suggested that alpha oscillations (~10 Hz) play a key role in this process. Increased alpha amplitude has been proposed to be associated with decreased excitability (for a review, see Foxe & Snyder, 2011; Jensen et al., 2012; Klimesch et al., 2007). In visual attention tasks, magnetoencephalography (MEG) and electroencephalography (EEG) studies have shown an increase in alpha amplitude, compared to baseline, over posterior regions ipsilateral to the attended side, particularly when a distractor was present on the unattended side (Green et al., 2017; Gutteling et al., 2021; Haegens et al., 2012; Ikkai et al., 2016; Kelly et al., 2006; Rihs et al., 2009; Sauseng et al., 2009; Siegel et al., 2008; Wildegger et al., 2017). In working memory tasks, alpha power increases have also been observed in the non-engaged stream (Bonnefond & Jensen, 2012a; Jokisch & Jensen, 2007; Magosso & Borra, 2024; Park et al., 2014; Payne et al., 2013; Tu et al., 2025), suggesting that alpha increases protect perceptual and working memory processes from distractors by decreasing neuronal gain in early visual regions.

Recently, several studies have questioned the link between alpha oscillations and functional inhibition (Antonov et al., 2020; Foster & Awh, 2019; Gundlach et al., 2020; Zhigalov & Jensen, 2020). These studies tested the relationship between excitability, as indexed by steady-state visual evoked potentials (SSVEPs), and alpha oscillations using EEG or MEG. They reported no clear link between non-attended SSVEPs and ipsilateral alpha increases in amplitude, space, or latency. These findings suggest that the relationship between alpha oscillations and functional inhibition requires further investigation. Furthermore, alpha modulation has primarily been reported in early visual regions, and whether a similar mechanism occurs in higher-order visual regions remains unclear (but see Capilla et al., 2014; Snyder & Foxe, 2010 for evidence of alpha modulation over the ventral stream). In a recent review paper, Bonnefond and Jensen (2025) challenged the prevailing paradigm by shifting the focus from whether an anticipatory alpha increase allows for optimal attention performance to the more fundamental question of when distractor suppression is actually necessary. They emphasized that distractor suppression is crucial when perceptual load is high (Lavie, 2005a), which is related to task difficulty, and/or high distractor interference, at perceptual or decision level. They further discussed the idea that when stimulus timing is predictable, alpha phase adjustment could be implemented as an alternative or additional mechanism as observed in a few attention and working memory tasks (Bonnefond & Jensen, 2012; Samaha & Postle, 2015; Solís-Vivanco et al., 2018 but see van Diepen et al., 2015). Finally, they suggest that inhibition of the ventral attention network (VAN), as observed in fMRI studies during goal-driven tasks (Corbetta & Shulman, 2002) and which is thought to prevent attentional capture may be sufficient, in particular when perceptual and decision interference is low. Our group was the first to demonstrate that VAN inhibition occurs via an increase in alpha/beta power during an attention task which correlated with resistance to the distractor interference (Solís-Vivanco et al., 2021).

The goal of the present study was to investigate (1) whether alpha oscillations are associated with functional inhibition in higher-order visual regions as well as in the VAN in a task involving high decision interference from distracting information and (2) whether alpha phase adjusts in anticipation of relevant and irrelevant stimuli.

To address these questions, we used MEG and designed a cue-based modified Stroop task inspired by the Stroop task (Stroop, 1935). In the Stroop task, participants report the ink color of a word that indicates a different color. The automatic reading of the word is known to interfere with the report of the ink color. In our version, a cue indicated whether participants should report the ink color (attend-color condition) or the color denoted by the word (attend-word condition). An fMRI study using the Stroop task reported increased BOLD signal in the “color area” and decreased activity in the visual word form area (VWFA) when participants reported the ink color (Polk et al., 2008). Given the negative correlation between alpha oscillations and BOLD signal observed in several EEG-fMRI studies (e.g. Laufs et al., 2006; Scheeringa et al., 2011), we hypothesized that alpha oscillations would increase in the VWFA and decrease in the color area in the attend-color condition. Furthermore, because our task design allowed participants to predict stimulus onset, we hypothesized that the alpha phase would be adjusted in anticipation of the stimuli, with distinct phases in the VWFA and color area during the attend-color condition. Specifically, we expected the phase adjusted in the VWFA to be associated with inhibition, e.g. as indexed by gamma decrease (Bonnefond & Jensen, 2015), to suppress word processing and the phase adjusted in the color area to be associated with excitation, to process the ink color optimally. We predicted that alpha amplitude and phase modulations would correlate with task performance. Finally, we hypothesized that alpha-beta activity (10-20 Hz) in the VAN would be high and associated with performance.

## Methods

### Participants

The study was carried out at the Donders Institute for Brain, Cognition and Behaviour. Twenty healthy native Dutch-speaking volunteers (age: 23 ± 2.95 years; 15 females) participated in the experiment after providing written informed consent according to the Declaration of Helsinki and the local ethics board. They were recruited from Radboud University’s research participation scheme. All subjects had normal or corrected-to-normal vision and were right-handed according to the Edinburgh Handedness Inventory (Oldfield, 1971).

### Stimulus material and procedure

The cue-based modified Stroop task (see **Figure 1**) was designed using MATLAB (MathWorks) and Psychtoolbox (psychtoolbox.org). The background of the screen was black during the entire experiment. Following an intertrial period of 1500 to 2000ms during which blinks were allowed, a trial started with a white dot presented during 1200ms (baseline period) followed by a cue during 100ms. Three different cues, representing abstract geometrical shapes, were presented (see **Figure 1**). One indicated to the participant to attend to the word (attend-word condition) while another one indicated to attend to the ink’s color (attend-color condition). These two cues were presented in 80% of trials. The third cue indicated that the participants would just have to determine whether a stimulus was presented in the trial (detect condition). Stimuli appeared in 80% of the trials in this condition. The association between a given cue and a given condition was randomized across participants. After a delay of 1200ms after cue offset (i.e., constant across trials), during which participants were asked to fixate the central white dot, visual stimuli were presented centrally for 100ms. The onset of the stimuli corresponded to time 0 in our analyses. We used three colors: red, blue, and yellow. Each word designating a color was associated with incongruent colors of the ink. The word blue (‘BLAUW’ in Dutch) was written in red or yellow, the word red (‘ROOD’ in Dutch) was written in blue or yellow, and the word yellow (‘GEEL’ in Dutch) was written in blue or red. Colored words were presented in 80% of the trials while no stimuli were presented in 20% of the remaining trials (525 trials in total). The participants also performed a similar task in fMRI another day (see procedure; data not analyzed here) with the aim of comparing Blood Oxygen Level Dependent (BOLD) signal and time-frequency power variations. Trials without stimuli (20% of the trials in each condition) were used in order to get a better BOLD signal estimate of the anticipatory activity in the fMRI data. They were also used in the present experiment in order to keep the design similar between recording methods. Participants were asked to respond as fast and accurately as possible by pressing one of the three buttons of a response pad. Each button was associated with a color and the association was randomized across participants. In the attend-color and attend-word conditions, participants were asked to press any button when no stimulus was presented. In the detect condition, the first and second buttons were associated with the ‘stimulus present’ and ‘stimulus absent’ responses respectively. Reaction time (RT) and response accuracy were recorded throughout the experiment. Trials of each condition were randomly presented.

**Figure 1.**
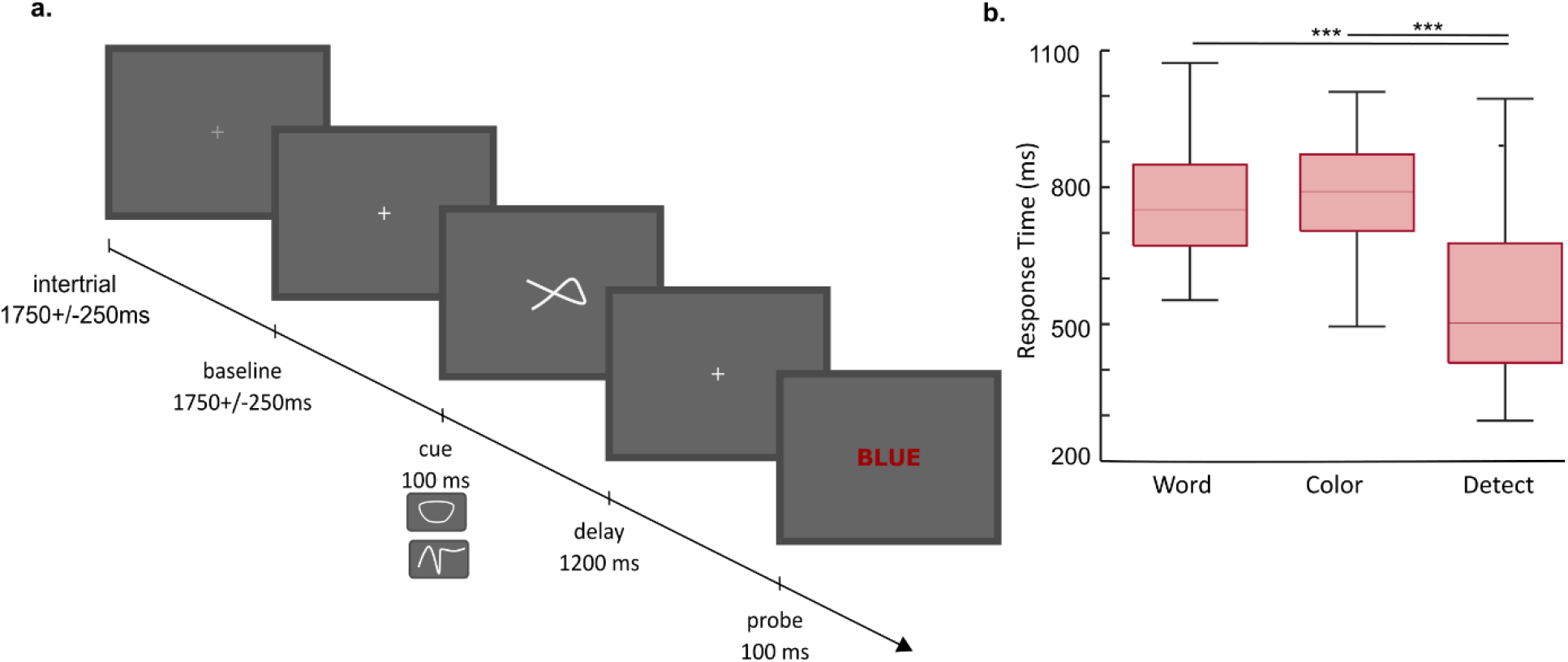
Task design and behavioral results. (**a**) Following a 1200ms baseline period, a cue (the three types of cue used are presented below the task design) instructed the participants to either attend the color of the ink (attend-color condition; 40% of the trials), the written word (i.e., the color designated by the word; attend-word condition; 40% of the trials) or to determine whether a stimulus was presented (detect condition; 20% of the trials). Participants were asked to respond as accurately and quickly as possible following the presentation of the stimuli. A stimulus was presented in 80% of the trials in each condition. (**b**) Behavioral results. Box plot of response times (RT) of all subjects for trials with correct answers and conditions with a stimulus presented on the screen. No significant difference between attend-word and attend-color conditions was observed. Significant differences between the median RT in word/color condition and detect condition were observed (*** = p<0.001).

In addition, two localizer tasks were performed. In the word localizer task, participants were presented with words, pseudo-words or checkerboards for 80ms with an intertrial interval of 800ms. We used 120 highly imaginable nouns (3-7 letters and 1-3 syllables in length). Pseudo-words were derived from them so that the frequency distribution of consonants was similar between words and pseudo-words. Checkerboards were rectangles and extended from 2^°^ to 6^°^ away from fixation with approximately the same vertical size as letter strings. The different categories of stimuli were presented in different blocks (6 per condition) of 20 trials. The task of participants was to detect deviants (words and pseudo-words with lower-case letters and checkerboards with missing black squares; 5% of trials). In the color localizer task, participants were presented with colored or grayscale paintings (215 trials in each condition). Stimuli were presented in the center of the screen and measured 10×6cm at a distance of 80cm. The task of the participants was to detect drawings of a child among known paintings (5% of trials). However, these tasks were performed at the end of the recording session, participants moved more than during the main task and were more tense, generating many muscular artifacts. An issue with the code of the triggers further impacted the analyses. The remaining number of trials did not allow us to localize the areas specialized in color and word processing in each participant and to perform region of interest analyses at the source level.

### Data acquisition

Neuromagnetic activity was recorded at a sampling rate of 1200Hz, using a 275 first-order axial gradiometers whole-head MEG system (VSF/CTF Systems, Port Coquitlam, Canada) housed in a magnetically shielded room. To measure the subject’s head position relative to the MEG during the experiment, three marker coils were placed respectively at the nasion and the left and right ear canals. During the experiment, horizontal and vertical eye movements were recorded using vertical and horizontal EOG electrodes. Subjects were in the supine position during the recording.

An anatomical T1 MRI of the participants was acquired with a 3T Siemens Sonata system (Erlangen, Germany) with a voxel size of 1mm^3^. For the co-registration of the MRI and the MEG data, tablets of Vitamin E were used and placed at the position of the coils.

### Procedure

The experiment was conducted over three consecutive days for each participant. During the first day, inclusion criteria were confirmed, general information about the study and informed consent letters were provided, and detailed instructions about the experiment were presented. Participants then performed a practice session composed of 150 trials inside the MEG room (60 trials for the two main conditions and 30 for the detect condition). This training session was necessary for the participants to get familiar with the cue-stimulus delay. This familiarity allowed us to test whether the alpha phase was adjusted in anticipation of stimuli (see Bonnefond & Jensen, 2012; Samaha et al., 2015; Solís-Vivanco et al., 2018).

During the second day, the MEG experiment or an fMRI experiment (preceded by a mock session within a dummy scanner) with a similar task was conducted (fMRI analyses performed separately). During the third day, the fMRI or the MEG experiment was performed depending on the neuroimaging technique used during the second session. During the fMRI session, the MRI of each subject was obtained. Participants were asked to wear no make-up during recordings and to wash their hair and change their clothes between the fMRI and the MEG session when the fMRI session occurred first. We did not observe different noise levels in the MEG recordings (using FieldTrip data quality check procedure) between participants who performed the fMRI first and those who performed the MEG first.

### Data analysis

*Hit rate* (HR; percentage of correct responses) was computed for each participant in each condition. *Reaction times* (RTs) were obtained from the subjects’ response pad responses. We computed the median of the RTs for each participant.

*The MEG analyses* were performed using the FieldTrip software package (Oostenveld et al., 2011). MEG data were epoched 1000ms before the onset of the cue until 600ms after stimulus onset. An automatic rejection, based on a z-score algorithm across sensors exceeding a threshold given by the data variance within each participant, of eye blinks or saccades, SQUID jumps, or muscle artifacts was run. Additional visual inspection was applied to the remaining trials before including them in further analyses. Only epochs without artifacts and with correct answers were considered. 80% of the trials were kept on average.

*For the sensor-level analyses*, planar gradients of the MEG field distribution were calculated (Bastiaansen & Knösche, 2000). For this purpose, we used a nearest neighbor method where the horizontal and vertical components of the estimated planar gradients were derived, therefore approximating the signal measured by MEG systems with planar gradiometers. This representation facilitates the interpretation of the sensor-level data, because the largest signal of the planar gradient is typically located above the source.

Time-frequency representations (TFRs) were obtained using a fast Fourier transformation approach with a 5-cycle long adaptive sliding time window (ΔT = 5/f; e.g., ΔT = 500 msec for the 10Hz frequency). A Hanning taper (ΔT long) was multiplied by the data before the Fourier transformation. For the planar gradient, the TFRs of power were estimated for the horizontal and vertical components and then summed. The power for the individual trials was averaged over conditions and log-transformed.

In order to determine the amplitude of the alpha activity phase-locked prior to stimulus onset, TFR of the power of averaged epochs (i.e., the event-related fields; ERF) were calculated as well.

*Source localization for the main task* was performed using a frequency domain beamforming approach based on an adaptive filtering technique (Dynamic Imaging of Coherent Sources, DICS; Gross et al., 2001). We obtained cross-spectral density matrices by applying a multitaper FFT approach (ΔT = 500 msec; one orthogonal Slepian taper) on data measured from the axial sensors. A realistically shaped single-shell description of the brain was constructed, based on the individual anatomical MRIs and head shapes (Nolte, 2003). The brain volume of each participant was divided into a grid with a 1-cm resolution and normalized to the template MNI brain (International Consortium for Brain Mapping, Montreal Neurological Institute, Canada) using SPM8 (www.fil.ion.ucl.ac.uk/spm). The lead field and the cross-spectral density were used to calculate a spatial filter for each grid point (Gross et al., 2001), and the spatial distribution of power was estimated for each condition in each participant. A common filter was used for both conditions, i.e., it was based on the cross-spectral density matrices of the combined conditions. The regularization parameter was set at 5%. We performed time-frequency analyses based on the results observed at the sensor level, and the estimated power was averaged over the time-frequency of interest and trials and log transformed. The contrast between conditions was performed and averaged across participants. The source estimates were plotted on a standard MNI brain found in SPM8.

*Region of interest (ROI) analyses* were performed using a time-domain beamformer, the linearly constrained minimum variance (LCMV) scalar beamformer spatial filter algorithm, to generate maps of source activity on a 1 cm grid (Van Veen et al., 1997).. The beamformer source reconstruction calculates a set of weights that maps the sensor data to time-series of single trials at the source locations, allowing to reconstruct the signal at the source level. In addition to TFR of power, we explored the correlation between power in different frequency bands and behavioral performances within and between subjects using Spearman’s rank correlations. We further investigated functional connectivity across these reconstructed time series by means of TFR of coherence (Rosenberg et al., 1989). We then tested for the directionality of the effects observed using non-parametric Granger causality (Dhamala et al., 2008). Finally, we performed cross-frequency coupling analysis using both the direct phase-amplitude coupling estimator developed by Özkurt and Schnitzler (2011) and the phase-locking approach developed by Cohen (2008). To perform these analyses, we adapted the PACmeg toolboxes (Seymour et al., 2017; https://neurofractal.github.io/PACmeg/).

### Statistics

Regarding the behavior, we tested for normality of the data using the Shapiro-Wilk test. Then, accuracy and log-transformed response time (RT) were analyzed using t-tests. We corrected for multiple comparisons using the Benjamini & Hochberg procedure for controlling the false discovery rate (FDR) of a family of hypothesis tests.

To establish statistical significance of power differences observed between conditions at both sensor and source levels, we used one-sided *t*-test computation. To control for the family-wise error rate (FWER), we employed a non-parametric cluster-based permutation test using Threshold-Free Cluster Enhancement (TFCE; *E* = 1, *H* = 2) as implemented in FieldTrip. The test was conducted across sensors, time and frequency or across the source space. Significance was determined by testing the actual TFCE score against the maximum TFCE scores yielded by the permutation distribution (1000 permutations, *P* < 0.05; Smith & Nichols, 2009).

In each ROI, we further used t-tests, after testing for normality of the data using the Shapiro-Wilk test, to compare correlation, connectivity measures and phase-amplitude coupling measures. We further corrected for multiple comparisons using the Benjamini & Hochberg procedure.

## Results

### Similar performances in the attend-color and in the attend-word condition

We did not observe differences between the attend-color and the attend-word and conditions in terms of accuracy (respectively: 92.48±5.19; 91.58±4.97; t(19)=−0.7, p=0.48, CI[−1.75 3.55]), but we did observe a significantly better accuracy in these conditions compared to the detect condition (97.3±3.6; attend-word vs detect: t(19)=−5.02, p=2e-4, CI[−6.83 −2.80]; attend-color vs detect: t(19)=−4.39, p=5e-4, CI[−8.45 −2.99]).

Analysis of RTs, for trials with correct answers, revealed no difference between the attend-color and the attend-word conditions although a trend was observed (see **Figure 1b**; respective median 750.8 ± 135.88 ms vs 790.4 ± 134.31 ms; t(19)=−1.43, p=0.16, CI[−43.85 8.23]). We observed a significantly faster reaction time in attend-word and attend-color conditions compared to the detect condition (median 502.1 ± 181.9 ms; t(19)= 7.95, p=1.83e-7 and t(19) =7.27, p=6.70e-7).

To summarize, subjects responded faster and with higher accuracy when they just had to detect whether a stimulus was presented. We did not observe a significantly slower RT in the attend-color condition compared to the attend-word condition, which may result from intensive training or from the short stimulus presentation duration.

We further analyzed the number of interference errors, i.e., reporting the color of the word in the attend-color condition and of the ink in the attend-word condition, with a repeated-measure ANOVA. We observed a higher number of interference errors compared to the other type of error (i.e., reporting the color that was not presented in the other dimension) in both conditions (F(1,19) = 25.16, p = 7.67×10^−5^).

### No significant difference in anticipatory alpha power and phase in attend-color vs attend-word condition

We then quantified the alpha power from the MEG data for the attend-color condition and the attend-word condition. The TFR of power did reveal several time-windows with increased alpha power in the attend-color condition over the left temporal sensors. These increases of alpha power occurred every~600ms with a burst occurring just before the onset of the stimulus. None of these effects did however survive multiple comparisons correction (p <0.05 before correction; see **Figure 2**).

**Figure 2.**
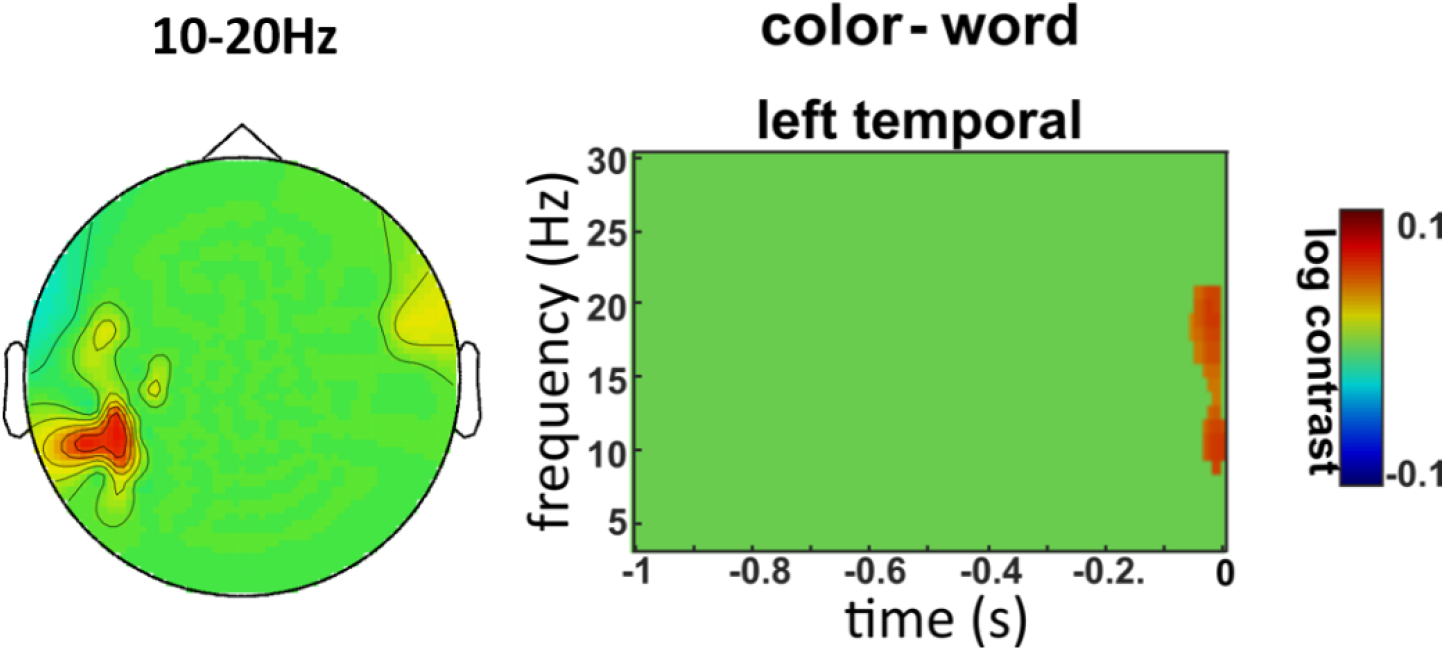
Alpha/beta power is stronger over the left temporal sensors. Results from the contrast between the color and the word conditions before stimulus onset and masked for significant results without correction for multiple comparisons. Time-frequency representations (TFR) of power from −1 to 0 s before the colored word onset show that 10-20Hz power was stronger in the attend-color condition, just before the stimulus onset, over left temporal sensors before correction for multiple comparisons.

Similarly, in the averaged epochs (testing for the alpha phase alignment), we found bursts of strong alpha power in the attend-color condition but, similarly to the analysis above, these effects did not survive multiple comparisons correction.

### Alpha/Beta and Theta increase in the ventral attention networks in anticipation of relevant stimuli

Following the approach we used in the study by Solis-Vivanco et al. (2021), we then compared the prestimulus period in word and color conditions combined to the pre-cue period (baseline) in these conditions. Following, the results observed in this study, we focused on power increase. We observed a power increase in the 4-6Hz and 10-20 Hz range over middle and right scalp regions before stimulus processing compared to pre-cue baseline ([attend-color+ attend-word]; **Figure 3a**). Statistics were applied in the −1 to 0s time window, in the 4-30Hz frequency range. We found significant differences between the two periods (TFCE-corrected for multiple comparisons; p < 0.001). The difference was driven by an effect from −0.7 to 0s within frequency ranges 4-6Hz and 10-20Hz over middle and right sensors.

**Figure 3:**
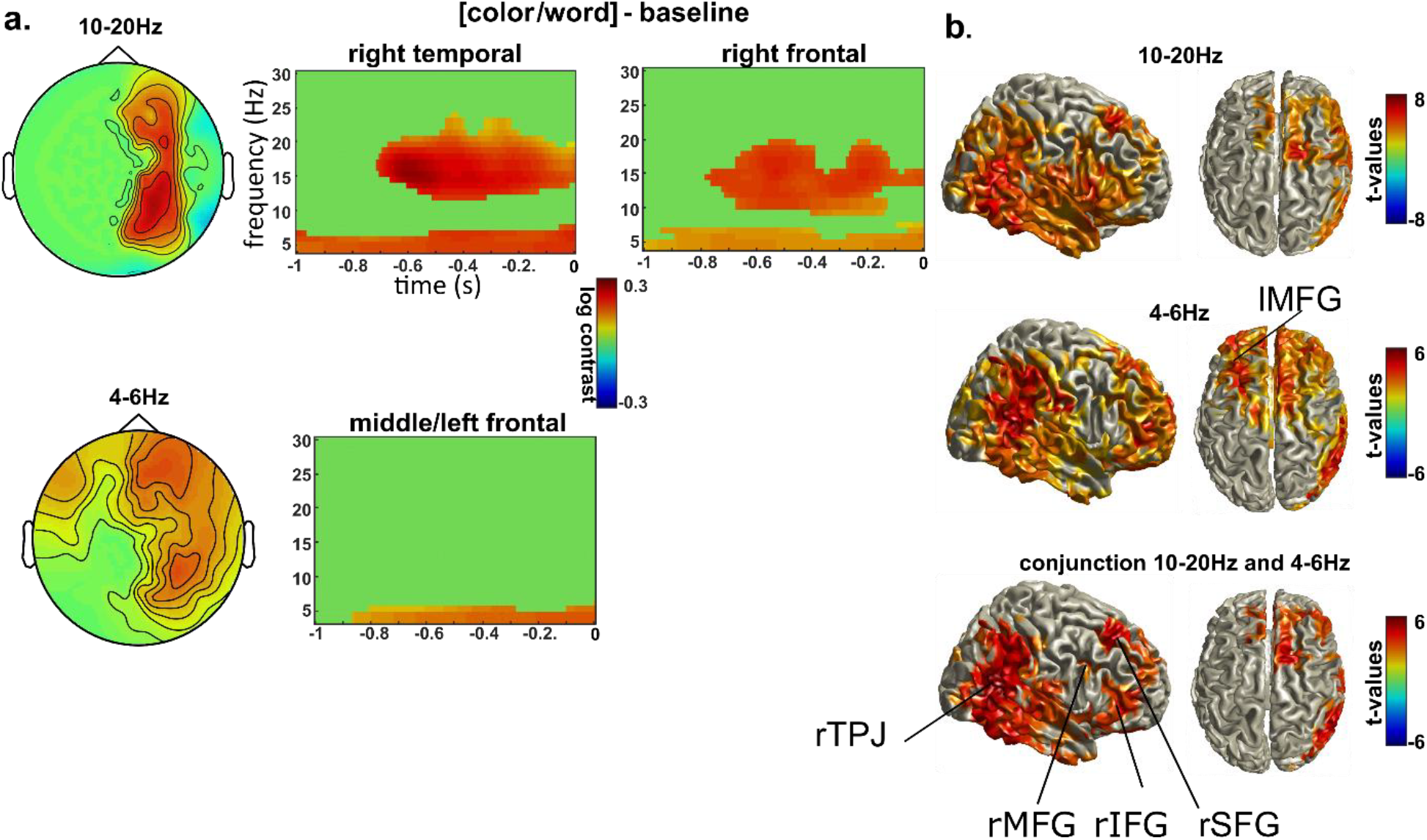
High alpha/beta and theta over the VAN and the left MFG. (a) Results from the contrast between pre-stimulus and pre-cue (baseline) period in the word and color conditions combined and masked for significant results only (TFCE corrected for multiple comparisons). Time-frequency representations (TFR) of power from −1 to 0 s before the colored word onset show that 4-6Hz and 10-20Hz power was stronger than during baseline over a right fronto-temporal cluster. 4-6Hz was further increased over left frontal sensors. (b) (top) Source localization of the significant right 10-20Hz increase compared to baseline (TFCE corrected). Most of these regions observed are part of the VAN. (middle) Source of the 4-6Hz increase. Regions observed are part of the VAN and of the CCN (bottom) conjunction of the two frequencies revealing the VAN network as well as the right and left SFG.

DICS beamformer was used to localize the sources of these effects. We used the AAL atlas by Tzourio-Mazoyer et al. (2002) to label the sources in which differences were observed. The significant difference (TFCE-corrected for multiple comparisons; p < 0.001) observed at the source level for the 15Hz (5Hz frequency smoothing; in 500ms time windows; −0.6 to −0.1s vs. −1.9 to −1.4s, i.e., pre cue) was driven by differences in the right temporo-parietal junction (rTPJ, MNI [58 −58 18]), the right inferior temporal gyrus (rITG; MNI [−70 −40 −22]), the right medial frontal gyrus (rMFG, MNI [42 20 50]), the right inferior frontal gyrus (rIFG; MNI [42 32 12]), the left and right dorsolateral superior frontal gyrus (l+rSFG; MNI [−16 38 42] and [20 30 52]) and the left and right anterior cingulate cortex (l+rACC; MNI [−2 34 22] and [4 24 26]). The rTPJ, the rIFG and the rMFG are nodes of the ventral attention network (VAN), the ACC is part of the cognitive control network (CCN) and the dorsolateral SFG is connected to these regions (see **Figure 3b**; top; Alves et al., 2019; Li et al., 2013).

The significant difference (TFCE-corrected for multiple comparisons; p < 0.001) observed at the source level for the 5Hz power band for the same time-windows (2Hz frequency smoothing) was also driven by differences within the same network (see **Figure 3b**; middle and bottom) as well as the left medial frontal gyrus (lMFG; MNI [−26 30 30]). This region has also been associated with the CCN (Cole et al., 2012; Cole & Schneider, 2007).

We also compared pre-stimulus period of word and color conditions combined to the same period in detect condition although the number of trials without artifacts in the latter condition was lower (N =81 vs N = 260). However, similar results as the ones observed for comparisons with pre-cue period were observed (results not shown here). In addition, the comparison to baseline for word and color conditions separated provided the same results as when they are combined (results not shown here).

### Alpha/Beta and Theta negatively correlated with reaction times in the ventral attention network

We then performed regions of interest (ROIs) analyses, using LCMV (see methods), in five regions in which differences were the strongest. The VAN was represented by the right IFG, the right MFG and the right TPJ and the CCN by the left MFG as it was further away from the other regions and exhibited theta activity only. We also included the right SFG which showed a strong effect. **Figure 4a** presents the TFR of the contrast of pre-stimulus and pre-cue periods in the different ROIs in word and color condition combined for illustrative purposes.

**Figure 4:**
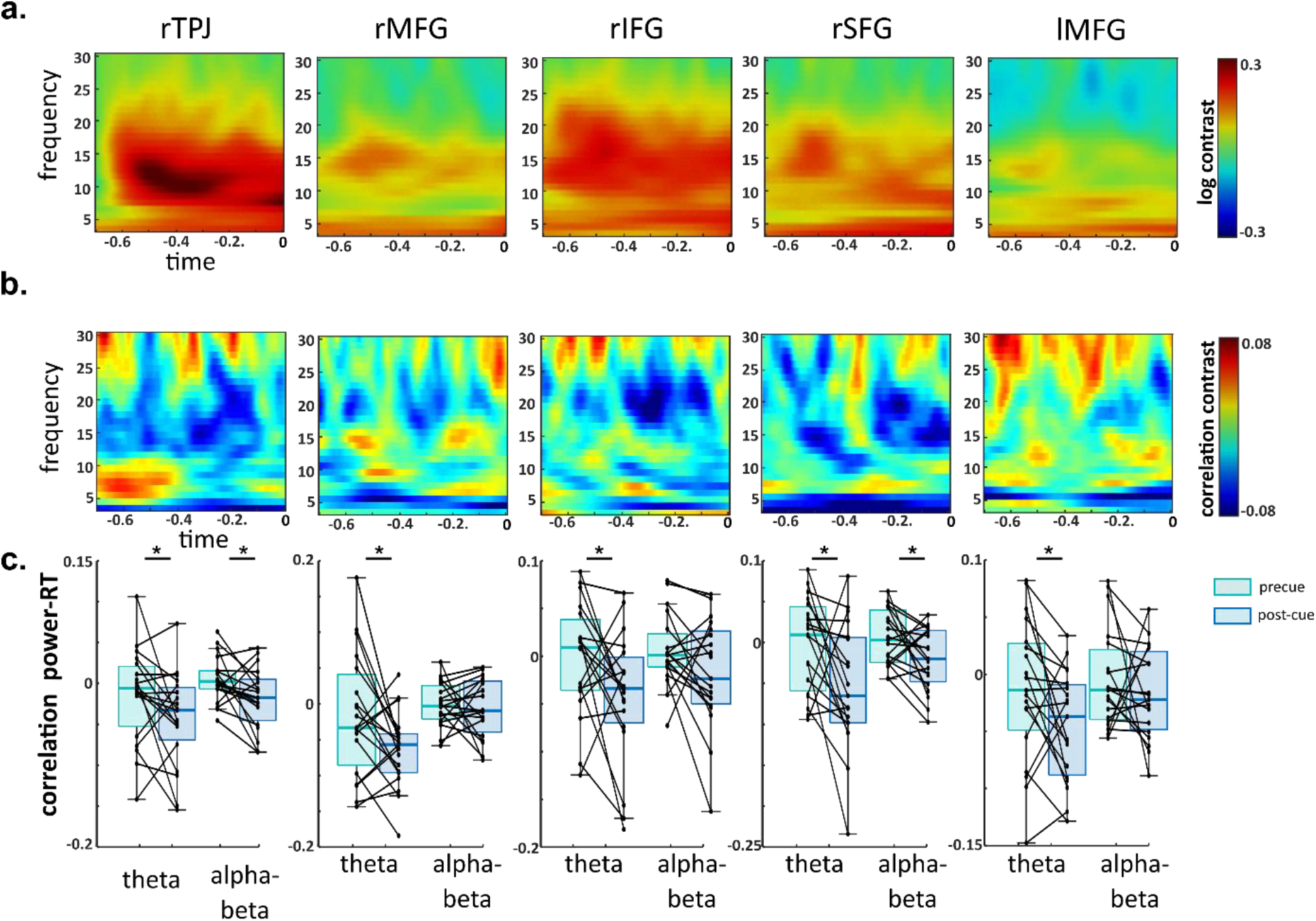
Theta power in the VAN and left MFG and beta power in right TPJ and right SFG are negatively correlated to reaction times. (a) TFR of the contrast between pre-stimulus and pre-cue (baseline) periods in the word and color conditions combined in the different ROIs. (b) Average TFR of the correlation between power and reaction times (RTs) normalized with the pre-cue period. (c) Box plots of the correlation with RTs of the averaged power in theta (4-6Hz) and alpha-beta (10-20Hz) frequency bands in the 250ms pre-stimulus period during pre-cue and pre-stimulus (post-cue) periods. Each dot represents a participant and each line connects the two periods. * indicates p < 0.05 after FDR correction for multiple comparisons.

We then aimed to determine whether these increases were associated with better performances over trials, i.e. whether there was a negative correlation between power and RTs. The Spearman correlation performed on this data over frequency and time with RTs revealed a negative correlation in theta frequency in all ROIs as well as alpha-beta frequency in several ROIs (see **Figure 4b**). To characterize these effects, we correlated log-transformed averaged power for theta (4-6Hz) and for alpha-beta (10-20Hz) in the 250ms pre-stimulus period with log-transformed reaction times and compared the correlation values obtained within a 250ms pre-cue period (see **Figure 4c**). These values were normally distributed (Shapiro-Wilk test, p > 0.05). One-sided t-tests revealed that post-cue theta was significantly correlated with RTs in all ROIs (p < 0.05 after Benjamini & Hochberg procedure; see **table 1** in supplementary information SI) while post-cue alpha-beta was correlated with RTs in the right TPJ and the right SFG (p < 0.05) although a trend was observed in the right IFG (p <0.05 before correction and p = 0.0789 after correction for multiple comparisons). Also, following the analysis performed in Solis-Vivanco et al. (2021), we explored the association between averaged power values (of both conditions combined) in each frequency band (and in each ROI) and the number of total interference errors (i.e., responses to the unattended feature compared to responses to the feature not displayed) over participants. We found that stronger power (baseline corrected) during the anticipatory period was inversely related with interference errors (r = −0.5279; p = 0.049 after correction for multiple comparisons; see **table 2** and supplementary **figure 1** in SI) in the beta frequency band in the right SFG. However, this kind of analysis should be considered with caution given the low number of participants (Schönbrodt & Perugini, 2013).

### The left medial frontal gyrus is synchronized with the right superior frontal gyrus and the right inferior frontal gyrus in the theta band

We aimed to determine whether the different nodes of the regions of interest were synchronized in the theta and/or alpha-beta frequency bands using coherence measures (see TFR of coherence in each pair of nodes in **Figure 5a**). To characterize these effects, we averaged coherence over theta (4-6Hz) and over alpha-beta (10-20Hz) in the 250ms pre-stimulus period and compared the values obtained to average coherence obtained within a 250ms pre-cue period (see inserts in **Figure 5a** and supplementary **figure 2**). One-sided t-tests revealed that theta coherence was significantly synchronized between the left MFG and the right IFG and SFG (p < 0.05 after Benjamini & Hochberg procedure). Many significant effects did not survive correction for multiple comparisons but exhibited a trend (see **table 3** in SI). We summarized the effects observed in **Figure 5b**

We also applied Granger causality measures to determine the direction of the significant theta synchronization in the same time window. Granger values were normally distributed (Shapiro-Wilk test, p > 0.05); we therefore applied a Wilcoxon signed rank test comparing the two directions. We observed stronger directionality from the right IFG to the left MFG than the opposite (p = 0.033) but no significant difference in directionality between the right SFG and the left MFG. We also attempted to characterize the relationship between theta phase and alpha-beta power, but again no robust pattern emerged when we compared approaches described in the methods section (results not shown here).

**Figure 5:**
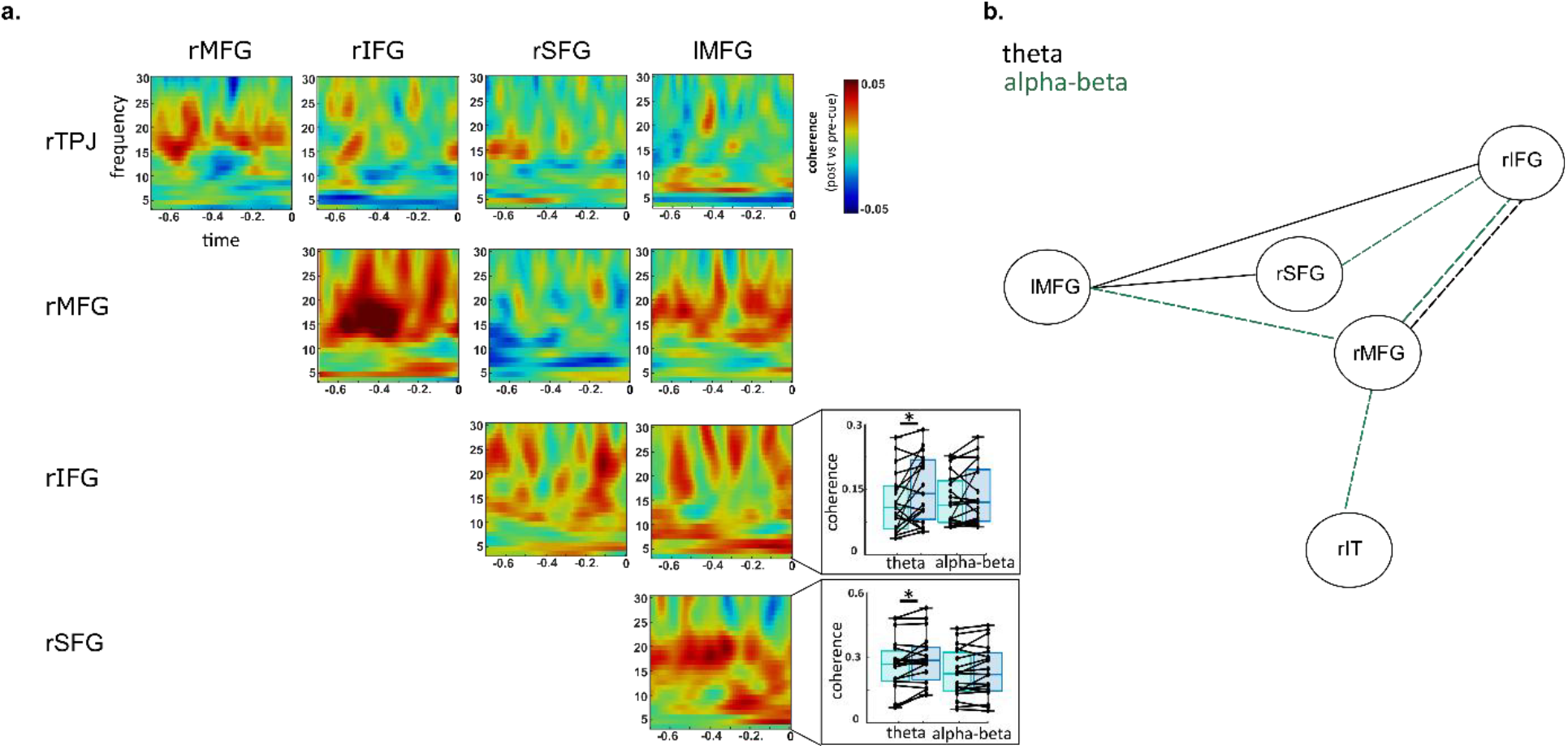
Theta synchronization between left MFG and right SFG and IFG during pre-stimulus period. (a) TFR of the coherence between each ROI during pre-stimulus compared to pre-cue (baseline) periods in the word and color conditions combined. (Right inserts) Box plots of the coherence between the lMFG and the rIFG and rSFG in theta (4-6Hz) and alpha-beta (10-20Hz) frequency bands during pre-cue and post-cue periods. Each dot represents a participant and each line connects the two periods. * indicates p < 0.05 after FDR correction for multiple comparisons. (b) Graphical representation of the significant effects that survived correction for multiple comparisons (full lines) and those that did not survive correction (dotted lines). Black lines represent theta coherence while green lines represent alpha-beta coherence.

## Discussion

In this study using a modified Stroop task, we had two main objectives. The first objective was to determine whether alpha oscillations are associated with functional inhibition over higher-order visual regions as well as over the ventral attention network (VAN; see Solís-Vivanco et al., 2021). The second objective was to determine whether alpha phase adjusts in anticipation of relevant and irrelevant stimuli. We did not find slower reaction times in the attend-color than in the attend-word condition, although a trend was observed. We found no significant modulation of alpha power and phase in anticipation of the stimuli in the attend-color compared to the attend-word condition. However, we found higher alpha-beta and theta power in the VAN and the left/right superior frontal gyrus (lSFG and rSFG) in anticipation of stimuli in both conditions. Theta power additionally increased in the left medial frontal gyrus (lMFG), a region associated with cognitive control. Theta power was negatively correlated with reaction times in all regions of interest (right temporo-parietal junction rTPJ, right medial frontal gyrus rMFG, right inferior frontal gyrus rIFG, rSFG and left medial frontal gyrus lMFG), while beta power was significantly negatively correlated with RTs in rTPJ and rSFG.

Behavioral results did not show the expected effect, i.e., response times slower in the attend-color than in the attend-word condition, as reading is automatic and difficult to overcome (Augustinova & Ferrand, 2014) and, as a consequence, words are expected to be particularly distracting when participants have to report the color of the ink (Stroop, 1935). These results could be explained by the long training of the participants, which is thought to increase the familiarity of participants with color naming and generally reduces the Stroop effect (Macleod, 1998; MacLeod & Dunbar, 1988).

As alpha power has been proposed to be associated with functional inhibition (Klimesch et al., 2007), we expected stronger pre-stimulus alpha power, and possibly alpha phase adjustment (see e.g. Solis-Vivanco et al., 2018), in the visual word form area (VWFA) in the attend-color compared to the attend-word condition. However, our time frequency analysis at the sensor level did not reveal a significant effect after correction for multiple comparisons. One explanation could be that some participants had a dominant VWFA over the right hemisphere, which could have blurred the group-level analyses. Indeed, a right-lateralization of the language network has been shown in ~7.5% of right-handed individuals and would be correlated with the lateralization of the VWFA (Gerrits et al., 2019; Knecht et al., 2000). Some studies have even revealed more variability for the lateralization of the VWFA than of the language network (see Carlos et al., 2019). Due to technical problems, we could not perform the region of interest analyses in the current task to test the distribution of the VWFA in our participants (see methods section). Another methodological limitation is the number of trials and participants in this study, as alpha increase has been shown to have a modest effect size that might require a large number of trials and participants (Wöstmann et al., 2022). This is in line with the trend we observed in the current data.

Beyond these possible methodological limitations, there is currently a hot debate in the literature regarding whether alpha oscillations are associated with the filtering of distracting information (Antonov et al., 2020; Foster & Awh, 2019; Zhigalov & Jensen, 2020). In a recent review paper, Bonnefond & Jensen (2025) emphasized in which contexts distractor filtering, and therefore alpha increase, could be implemented. Among the factors discussed, the most relevant here are the difficulty of the relevant task, more precisely the perceptual load, and the decision interference level of distractors. According to Lavie (2005), participants would particularly need to efficiently filter out distractors when the perceptual load is high. Gutteling et al. (2021) tested this hypothesis by varying the noise level of faces to manipulate the perceptual load of the target stimuli and the salience of the distractors. They found stronger ipsilateral alpha power (i.e., contralateral to the distractor) in the high load condition than in the low load condition. It is therefore possible that the perceptual load and/or decision interference of the distracting information, e.g., due to training and the associated increase in familiarity with color naming, were too low in the current task to induce a necessity for distractor filtering and hence an increase in alpha oscillations over the VWFA.

We found an increase in pre-stimulus alpha-beta in several nodes of the VAN in the attend-word and attend-color conditions compared to baseline, and a control condition, and a link to behavioral performance. This network has been associated with the capture of attention by unattended, but relevant, stimuli (Corbetta et al., 2008; Corbetta & Shulman, 2002; ElShafei et al., 2020). Corbetta et al. (2002, Corbetta et al. 2008) suggested that the inhibition of the VAN, as indexed by a decrease in the BOLD signal in fMRI, was crucial for preventing the capture of attention by distracting stimuli. In line with that idea, Solís-Vivanco et al. (2021) reported that higher alpha-beta power in the VAN was associated with lower distractor interference across subjects. Although the negative correlation with RT could be interpreted along this line, we did not observe a significant correlation with interference in these nodes. This might result from the low number of participants (Schönbrodt & Perugini, 2013) but also from the low number of errors, including interferences per participant, probably due to training.

In addition, we observed higher alpha-beta over the superior frontal gyrus (SFG). Alpha-beta power in this region was negatively correlated with RTs within participants and with interference errors across participants (but see Schönbrodt & Perugini, 2013). This region is not considered as being part of the VAN but has been reported to be connected to this network (Li et al., 2013). We previously observed this region in Solís-Vivanco et al. (2021). Li et al. (2013) proposed that this region is involved in the execution of cognitive manipulations, although its exact mechanistic role is unclear and further investigations are required.

Theta power increased in the anterior cingulate cortex (ACC) and the left medial frontal gyrus (lMFG), which are nodes of the cognitive control network (CCN; Matsumoto & Tanaka, 2004) and have been observed in Stroop tasks using fMRI (Polk et al., 2008; Roelofs et al., 2006). Some studies revealed a correlation between theta power over midfrontal and dorsolateral regions and conflict effects (e.g. Cohen & Donner, 2013). A recent study revealed that the anticipation of conflict was associated with increased frontal theta and that this measure was associated with faster conflict resolution (Kaiser et al., 2022). This is in line with our data. More surprising was the theta observed over the VAN and the SFG. Theta power was not analyzed in Solís-Vivanco et al. (2021), but Figure 2 in that paper indicates an increase in theta power over similar sensors as the beta increase. We hypothesized that the cognitive control network, represented by the left MFG, would control the VAN activity through synchronization in the theta band. Connectivity results indicated that the left MFG was synchronized with the right IFG and the right SFG. In addition, a trend was observed between the right IFG and the right MFG. A trend was further observed in the beta band between the nodes of the VAN, similarly to the results observed in Solis-Vivanco et al. (2021), and between the right IFG and the right SFG. We hypothesize that the left MFG controls the VAN through the right IFG region and indirectly through the right MFG region. However, the directionality measure indicates that the right IFG leads the left MFG, contradicting that hypothesis. Additional experiments would be required to confirm this result. Additional connectivity could occur in the alpha-beta band with the VAN. We further performed cross-frequency coupling to determine whether beta power was nested with the theta cycle in the VAN with the idea that the CCN controlled the VAN activity in the theta band, which locally controlled the beta band in the different regions of the VAN. However, the different measures used did not provide conclusive results and further investigation is required.

To conclude, we showed that both beta power over the VAN and theta power over the CCN are important to perform well in a modified Stroop task. We further showed that these two networks are connected in the theta band.

## Acknowledgments

The authors thank Ole Jensen for his help on designing the task and his comments on the manuscript. The authors also thank Rocio Silva, Jessica Askamp, and Paul Gaalman for their technical assistance. This work was supported by the European Research Council under the European Union’s Seventh Framework Programme (FP7/2007–2013)/ERC starting grant (grant number 716862) attributed to Mathilde Bonnefond. This work was conducted in the framework of the LabEx Cortex (“Construction, Function and Cognitive Function and Rehabilitation of the Cortex,” ANR-10-LABX-0042) of Université de Lyon.

