## Supplementary data for "Alpha-Beta oscillations implement inhibition of the ventral attention network during an attention task"

**Supplementary information**

| | | rTPJ | rMFG | rIFG | rSFG | lMFG |
|---|---|---|---|---|---|---|
| **Theta correlation** | Tstat(18) | -1.9945 | -2.4074 | -2.0822 | -2.4760 - | 2.1407 |
| | adjusted p value | 0.0307 | 0.0307 | 0.0307 | 0.0307 | 0.0307 |
| | CI | [-∞ -0.0038] | [-∞ -0.0134] | [-∞ -0.0065] | [-∞ -0.0155] | [-∞ -0.0059] |
| **Alpha Beta correlation** | Tstat(18) | -2.3487 | -0.4644 | -1.7638 | -2.3923 | -0.8438 |
| | adjusted p value | 0.0381 | 0.3240 | 0.0789 | 0.0381 | 0.2562 |
| | CI | [-∞ -0.006] | [-∞ -0.00153] | [-∞ -0.0004] | [-∞ -0.0072] | [-∞ -0.0128] |

***Table 1*: Statistics of the correlation with reaction times in different frequency bands**

| | | rTPJ | rMFG | rIFG | rSFG | lMFG |
|---|---|---|---|---|---|---|
| **Theta correlation** | r | -0.0720 - | -0.0552 | -0.2171 | 0.0578 | 0.1961 |
| | adjusted p value | 0.5139 | 0.5139 | 0.5139 | 0.5139 | 0.7894 |
| **Alpha Beta correlation** | r | -0.1933 | 0.1086 | -0.1007 | -0.5279 | 0.0633 |
| | adjusted p value | 0.5349 | 0.6709 | 0.5681 | 0.0499 | 0.6709 |

***Table 2*: Statistics of the correlation with interference errors over participants in different frequency bands**

| | | | rMFG | rIFG | rSFG | lMFG |
|---|---|---|---|---|---|---|
| **Theta coherence** | **rTPJ** | Tstat(18) | -1.3796 | -1.2806 | -0.7193 | -2.6658 |
| | | adjusted p value | 0.9921 | 0.9921 | 0.8169 | 0.9921 |
| | | CI | [-0.0347 ∞] | [-0.0239 ∞] | [-0.0215 ∞] | [-0.0321 ∞] |
| | **rMFG** | Tstat(18) | | 2.0358 | 0.0254 | 1.5383 |
| | | adjusted p value | | 0.0946 | 0.8167 | 0.1767 |
| | | CI | | [0.0037 ∞] | [-0.0214 ∞] | [-0.0019 ∞] |
| | **rIFG** | Tstat(18) | | | 1.2544 | 2.4659 |
| | | adjusted p value | | | 0.2258 | 0.0499 |
| | | CI | | | [-0.0059 ∞] | [0.09 ∞] |
| | **rSFG** | Tstat(18) | | | | 3.0558 |
| | | adjusted p value | | | | 0.034 |
| | | CI | | | | [0.0115 ∞] |
| **Apha-Beta coherence** | **rTPJ** | Tstat(18) | 2.2221 | -0.5555 | 0.0601 | -1.2742 |
| | | adjusted p value | 0.0656 | 0.8841 | 0.6805 | 0.8906 |
| | | CI | [0.0027 ∞] | [-0.0154 ∞] | [-0.0076 ∞] | [-0.0143 ∞] |
| | **rMFG** | Tstat(18) | | 2.3263 | -1.1094 | 2.4179 |
| | | adjusted p value | | 0.0656 | 0.8906 | 0.0656 |
| | | CI | | [0.0059 ∞] | [--0.0168 ∞] | [0.0062 ∞] |
| | **rIFG** | Tstat(18) | | | 1.8378 | 1.2242 |
| | | adjusted p value | | | 0.1033 | 0.1831 |
| | | CI | | | [0.0011 ∞] | [-0.0044 ∞] |
| | **rSFG** | Tstat(18) | | | | 1.7013 |
| | | adjusted p value | | | | 0.1061 |
| | | CI | | | | [-0.0002 ∞] |

***Table 3*: Statistics of the coherence between regions of interest in different frequency bands**

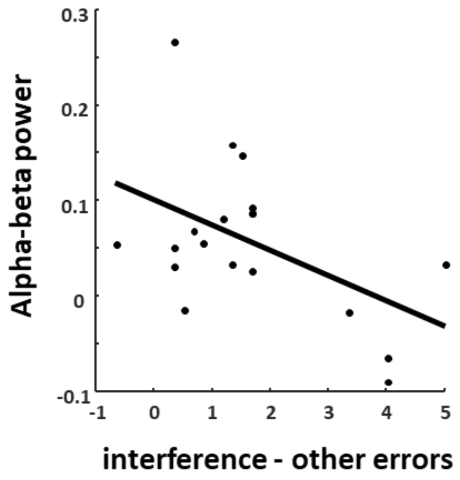


***Supplementary figure 1.*** Correlation between baseline corrected alpha-beta power and interference errors in the left superior frontal gyrus

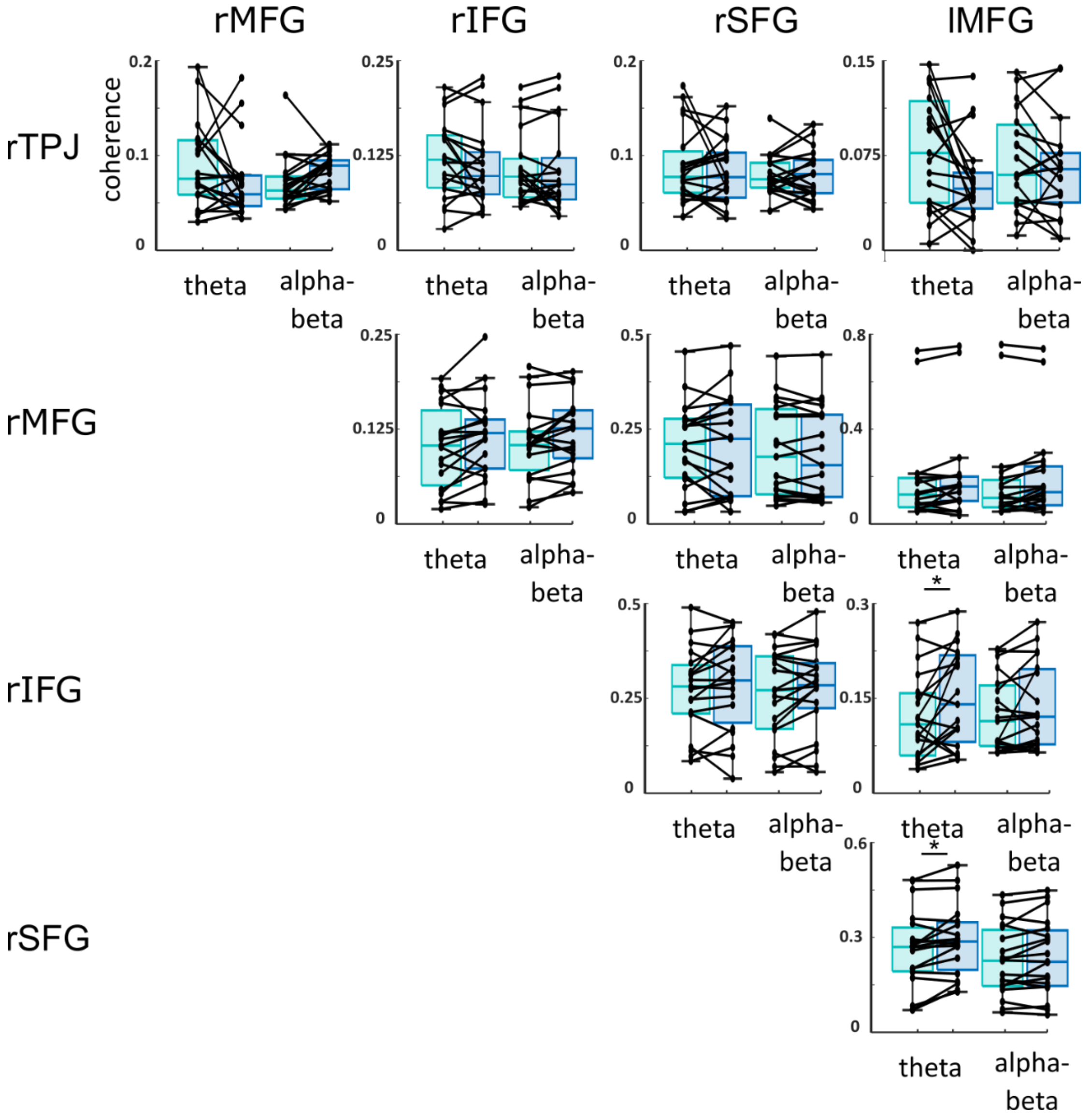


***Supplementary figure 2*** Plots of coherence measures between all regions of interest in the theta and alpha-beta frequency range***.***
